# Screening thousands of transcribed coding and non-coding regions reveals sequence determinants of RNA polymerase II elongation potential

**DOI:** 10.1101/2021.06.01.446655

**Authors:** Hanneke Vlaming, Claudia A. Mimoso, Benjamin JE Martin, Andrew R Field, Karen Adelman

## Abstract

Organismal growth and development rely on RNA Polymerase II (RNAPII) synthesizing the appropriate repertoire of messenger RNAs (mRNAs) from protein-coding genes. Productive elongation of full-length transcripts is essential for mRNA function, however what determines whether an engaged RNAPII molecule will terminate prematurely or transcribe processively remains poorly understood. Notably, despite a common process for transcription initiation across RNAPII-synthesized RNAs^1^, RNAPII is highly susceptible to termination when transcribing non-coding RNAs such as upstream antisense RNAs (uaRNAs) and enhancers RNAs (eRNAs)^2^, suggesting that differences arise during RNAPII elongation. To investigate the impact of transcribed sequence on elongation potential, we developed a method to screen the effects of thousands of <u>IN</u>tegrated <u>S</u>equences on <u>E</u>xpression of <u>R</u>NA and <u>T</u>ranslation using high-throughput sequencing (INSERT-seq). We found that higher AT content in uaRNAs and eRNAs, rather than specific sequence motifs, underlies the propensity for RNAPII termination on these transcripts. Further, we demonstrate that 5’ splice sites exert both splicing-dependent and autonomous, splicing-independent stimulation of transcription, even in the absence of polyadenylation signals. Together, our results reveal a potent role for transcribed sequence in dictating gene output at mRNA and non-coding RNA loci, and demonstrate the power of INSERT-seq towards illuminating these contributions.

## Introduction

The variability in RNAPII behavior is striking at divergent mammalian promoters, where an mRNA and uaRNA are transcribed in close proximity to one another, initiating from indistinguishable sequence elements, yet have vastly different fates^1^. Whereas mRNAs are long, precisely processed and stable, uaRNAs are generally <2kb in length, unspliced, unstable and non-polyadenylated^2^. This dichotomy suggests that differences arise during RNA elongation and lie within the transcribed sequences themselves.

While our understanding of RNAPII transcription regulation by sequence content is incomplete, some important sequence elements have been described. Most well-known is the polyadenylation signal (PAS), which consist of a PAS hexamer (canonically AAUAAA, but variants exist^3^) and several auxiliary elements that are recognized by the cleavage and polyadenylation (CPA) machinery^4^. PASs mediate the termination of most full-length mRNAs, and influence some fraction of premature termination in mRNAs and non-coding RNAs as well^5–8^. On the other hand, the process of splicing has long been suggested to stimulate transcription, although the mechanisms underlying this process remain unclear. Various reports implicate splicing in stimulation of transcription initiation, pause-release, or through the suppression of premature termination^9–14^. Notably, a splicing-independent role for the 5’ splice site (5’SS) has been suggested^6,8,10,15–17^, and the U1 snRNP can bind to both a 5’SS and RNAPII without the full spliceosome present^18^. However, prior experimental approaches to dissect the function(s) of 5’ SSs have suffered several limitations, including reliance on a small number of introns under study^10–12,17^ or the use of global splicing inhibitors^6,8,13,15,16^, which cause cellular toxicity and lead to indirect or off-target effects. Moreover, most analyses have focused narrowly on whether the presence of a 5’SS suppresses premature termination at cryptic PASs, in a process called ‘telescripting’^15,16^, leaving open the question of a more general role in stimulating transcription. Thus, whether RNA splicing and/or 5’SSs generally promote transcription remains to be systematically evaluated.

## Results

We developed INSERT-seq to decipher how transcribed sequence affects gene expression, and to answer open questions in the field regarding the role of splice sites in stimulating transcription. A fluorescent reporter sequence was integrated at a non-coding uaRNA locus in mouse embryonic stem cells (mESCs, Fig. 1a), and a library of thousands of sequences was introduced specifically at this allele, between the transcription start site (TSS) and reporter (Fig. 1b). The library comprised a repertoire of sequences from coding and non-coding transcripts, and synthetic sequences, designed to fully explore the relationship between sequence composition and expression.

**Figure 1.**
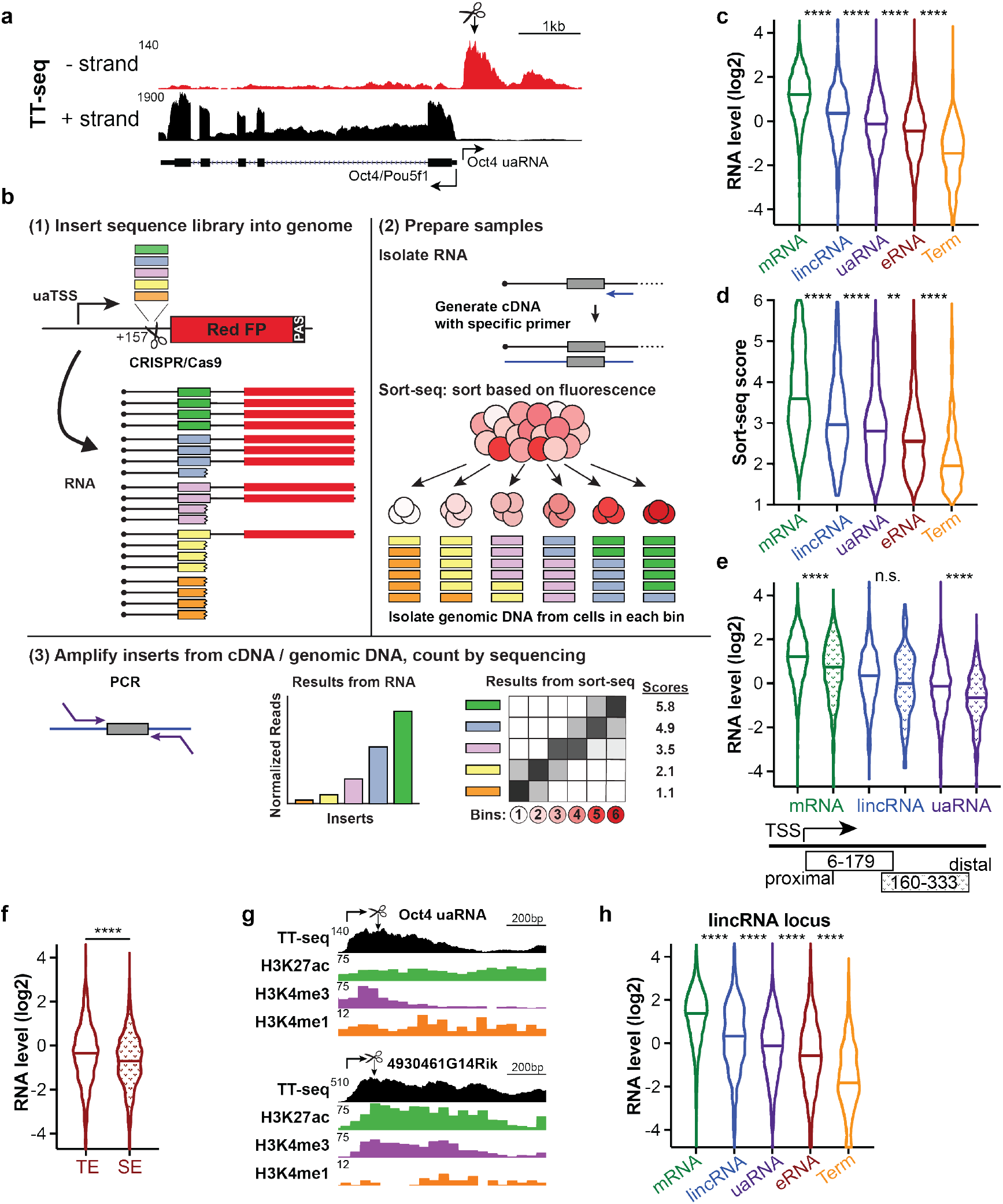
INSERT-seq demonstrates role of transcribed sequences in gene regulation. **a**, TT-seq data at the *Oct4* locus in mESCs, measuring newly synthesized RNA. As described in methods, a red fluorescent reporter was integrated into one allele, 172bp from the TSS, as indicated by arrow and scissors. **b**, Schematic of INSERT-seq. At the reporter locus, a library of inserts with 173bp variable regions was introduced by CRISPR/Cas9. To measure the effects of each insert on RNA abundance, RNA (steady-state or nascent) was isolated from the pool of cells, inserts were amplified from cDNA (made with gene-specific primer) and counted by next-generation sequencing. Abundance of inserts in RNA was normalized to abundance in genomic DNA. Protein expression was measured by sorting cells based on their level of red fluorescence (see methods) and the abundance of inserts in genomic DNA from each fluorescence bin used to calculate a Sort-seq score. **c,d**, Steady-state RNA (c) and Sort-seq (protein; d) results of inserts containing TSS-proximal genomic regions and mRNA terminators. Violins show frequency distribution and median. mRNAs *n* = 3,867, lincRNAs *n* = 342, uaRNAs *n* = 1,743, eRNAs *n* = 2,106, mRNA terminators *n* = 416. Comparisons between neighbors by Kruskal Wallis test. **e**, Steady-state RNA levels of inserts containing TSS-proximal (same as panel c) and TSS-distal genomic regions of indicated RNA classes. For TSS-distal regions, mRNAs *n* = 944, lincRNAs *n* = 98, uaRNAs *n* = 557. Comparisons by Kruskal-Wallis test. **f**, Steady-state RNA levels of inserts containing TSS-proximal regions from typical enhancers (TE, not overlapping super enhancers, *n* = 1,506) and super enhancers (SE, defined by ref.^21^, *n* = 600), compared by Mann-Whitney test. **g**, Snapshots of TT-seq and ChIP-seq data at the *Oct4* uaRNA and *4930461G14Rik* lincRNA loci. See methods for data sources. Transcription start sites and cut/integration sites are indicated; at the lincRNA locus the library was integrated 102bp downstream of the TSS. **h**, Steady-state RNA levels after library integration at the *4930461G14Rik* lincRNA locus, showing the same inserts classes as in panel c. mRNAs *n* = 3,987, lincRNAs *n* = 381, uaRNAs *n* = 1,853, eRNAs RNA terminators *n* = 458. All comparisons between neighbors are significant by Kruskal-Wallis test. For all statistical tests in this figure: n.s. *P* > 0.05, \*\**P* < 0.01, \*\*\*\**P* < 0.0001.

Effects of inserted sequences on RNA and protein levels were read out in high-throughput assays (Fig. 1b). In RNA assays, the relative enrichment of each insert in cDNA was determined, as compared to genomic DNA. cDNA was generated using a reverse primer that anneals 221nt downstream of the inserted sequences, such that insert abundance reflects the number of transcripts elongated past this location. Importantly, both steady-state RNA and nascent RNA (described below, Fig. 2c) can be used as starting material for this assay. Effects on reporter protein expression were measured using Sort-seq^19,20^: the pool of cells was sorted into six bins based on the level of red fluorescence relative to a control fluorophore (see Methods) and the inserts present in genomic DNA from each bin were sequenced. For each insert, the distribution over the six bins was used to calculate a Sort-seq score ranging from 1 (low) to 6 (high) protein abundance. Both screening methods showed good agreement between biological replicates (Extended Data Fig. 1a).

**Figure 2.**
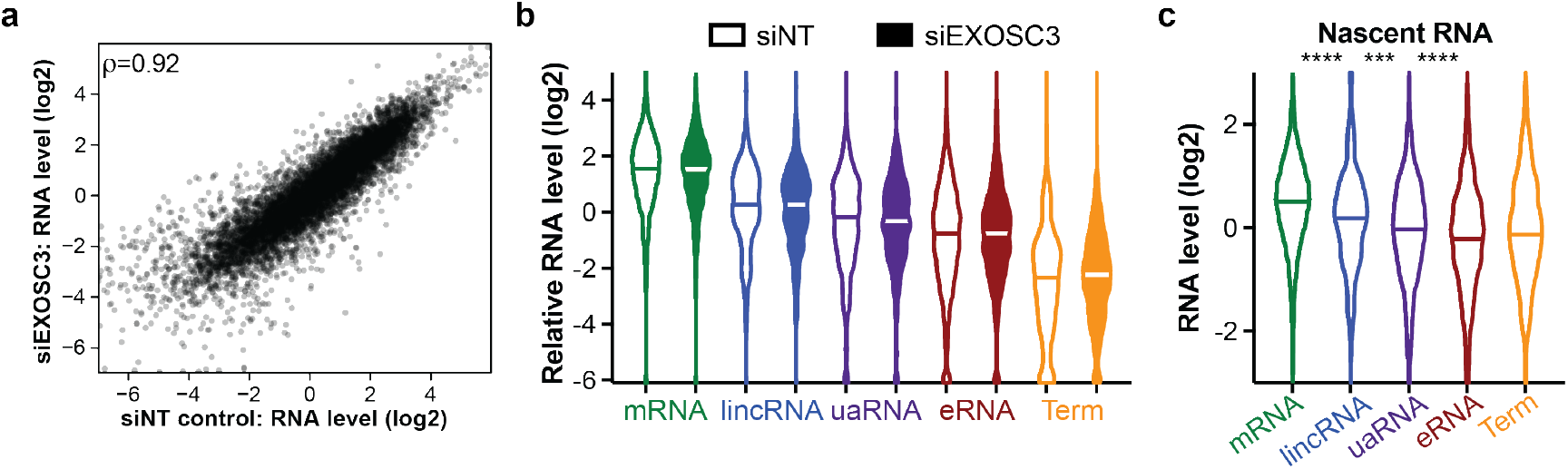
Transcribed sequence directly affects transcription levels. **a**, Correlation between relative abundance of inserts in steady-state RNA from cells 48 hours after transfection with a nontargeting control siRNA (siNT) or siRNAs targeting the exosome subunit EXOSC3. Plotted are all the inserts used for Fig. 1, each data point is the mean of three replicate experiments. **b**, Steady-state RNA results with library at uaRNA locus, comparing control siRNA (siNT, open) to exosome depletion (siEXOSC3, filled). mRNAs *n* = 3,356, lincRNAs *n* = 304, uaRNAs *n* = 1,480, eRNAs *n* = 1,749, mRNA terminators *n* = 333. **c**, Nascent RNA results with library at uaRNA locus. Nascent RNA was isolated after biotin-rNTP run-on (methods). Groups as in in Fig. 1c. Neighbors were compared by Kruskal-Wallis test. \*\*\**P* < 0.001, \*\*\*\**P* < 0.0001.

### 5’ end sequence contributes to transcript expression

First, we asked whether the composition of the transcribed sequence affects its own transcription. To this end, the library included TSS-proximal regions from different transcript classes. As a negative control, mRNA termination regions were included, which exhibited low abundance (Fig. 1c,d), as expected. TSS-proximal regions from mRNAs supported the highest levels of RNA and protein, followed by long intergenic non-coding RNAs (lincRNAs), and then uaRNAs and eRNAs (Fig. 1c,d). This is consistent with the endogenous elongation potential for these transcript classes and suggests that the composition of the TSS-proximal sequence contributes to transcription regulation. Next, we tested whether functional sequence elements were limited to the most TSS-proximal regions or could be found equally in TSS-distal regions (roughly 160-333bp from the TSS) (Fig. 1e, Extended Data Fig. 1b). Interestingly, TSS-proximal regions supported higher abundance than their distal counterparts. Thus, we suggest that the mammalian genome has evolved to contain signals within the initially transcribed region that directly influence elongation potential.

We specifically evaluated sequences from typical enhancers (TE) and super enhancers (SE, ref. ^21^), since it has been proposed that transcription termination is particularly important within SEs to prevent collisions between closely spaced, convergently transcribing RNAPII^22^, which could cause DNA damage. In agreement with this idea, we observed that eRNA sequences from SEs are consistently more negative than those in TEs (Fig. 1f, Extended Data Fig. 1c), suggesting that eRNA sequences themselves contribute to efficient termination observed in SEs.

### Effect of the transcribed sequence is independent of context

To test whether these conclusions were generalizable, we repeated the RNA-level screen after integrating the sequence library downstream of the *4930461G14Rik* lincRNA TSS. We selected this locus, since this lincRNA is many kb long, highly transcribed and, in comparison to the uaRNA locus, displays higher levels of H3K27ac, a larger domain of H3K4me3 and less H3K4me1 (Fig. 1g). Screening the library at this lincRNA locus, without introduction of a fluorescent reporter, revealed a strong correlation with the data obtained at the uaRNA locus (Fig. 1h, Extended Data Fig. 1d). We conclude that the signals contained in transcribed sequences can be conveyed independent of chromatin context or promoter identity.

### Transcribed sequence directly affects transcription levels

Since uaRNAs and eRNAs have been reported to be unstable due to rapid degradation by the exosome complex^2,23–25^, we tested whether the differences in RNA abundance observed could be explained by RNA stability. First, we repeated the RNA screen with the library at the uaRNA locus after knockdown of the exosome subunit EXOSC3 (RRP40; Extended Data Fig. 2a). While EXOSC3 depletion resulted in a general stabilization of transcripts from the uaRNA reporter locus (Extended Data Fig. 2b), the relative effects of inserts were the same as in the control condition (Fig. 2a), with mRNAs regions remaining the most positive and eRNAs the most negative (Fig. 2b). Second, we isolated nascent RNA from the library-containing pool of cells to use as the starting material for an RNA-based screen. To this end, nascent RNA was run on with biotinylated rNTPs, and labeled RNAs were purified using a modified version of PRO-seq^26^. Consistently, we found the pattern of mRNA>lincRNA>uaRNA>eRNA in insert abundance in the nascent RNA (Fig. 2c), and a good correlation between nascent and steady state RNA levels (Extended Data Fig. 2c). Taken together, these results show that the transcribed sequence can directly affect the levels of its own transcription.

### GC content inherently affects transcriptional output

mRNAs are more GC-rich than the average genomic region, especially at 5’ ends, which often overlap with CpG islands^1^. In contrast, the GC content of non-coding RNAs are closer to genomic background level of 42% G/C in mice^27^ (Extended Data Fig. 3a). To test whether GC content influences expression levels, we included in our library ~1000 synthetic control sequences generated over a range of GC contents that paralleled the genomic inserts in the library. Notably, these synthetic controls were designed to be devoid of the two most common PAS hexamers, to remove canonical termination sequences. Evaluation of the control sequences demonstrated that inserts with higher GC content had a higher abundance in all assays (Fig. 3a, Extended Data Fig. 3b,c), indicating a striking impact of GC content on RNA production. To determine if it is high GC content, or a specific enrichment in CpG dinucleotides at 5’ ends that promotes transcription^28^, we compared these features with RNA abundance of control sequences. These analyses demonstrated that CpG content is not a better predictor of RNA level than overall GC content (Pearson correlation coefficient of 0.47 in both cases, Fig. 3a and Extended Data Fig. 3d). We therefore conclude that early-transcribed regions that are enriched in G and C nucleotides support higher levels of transcription than sequences with a high AT content.

**Figure 3.**
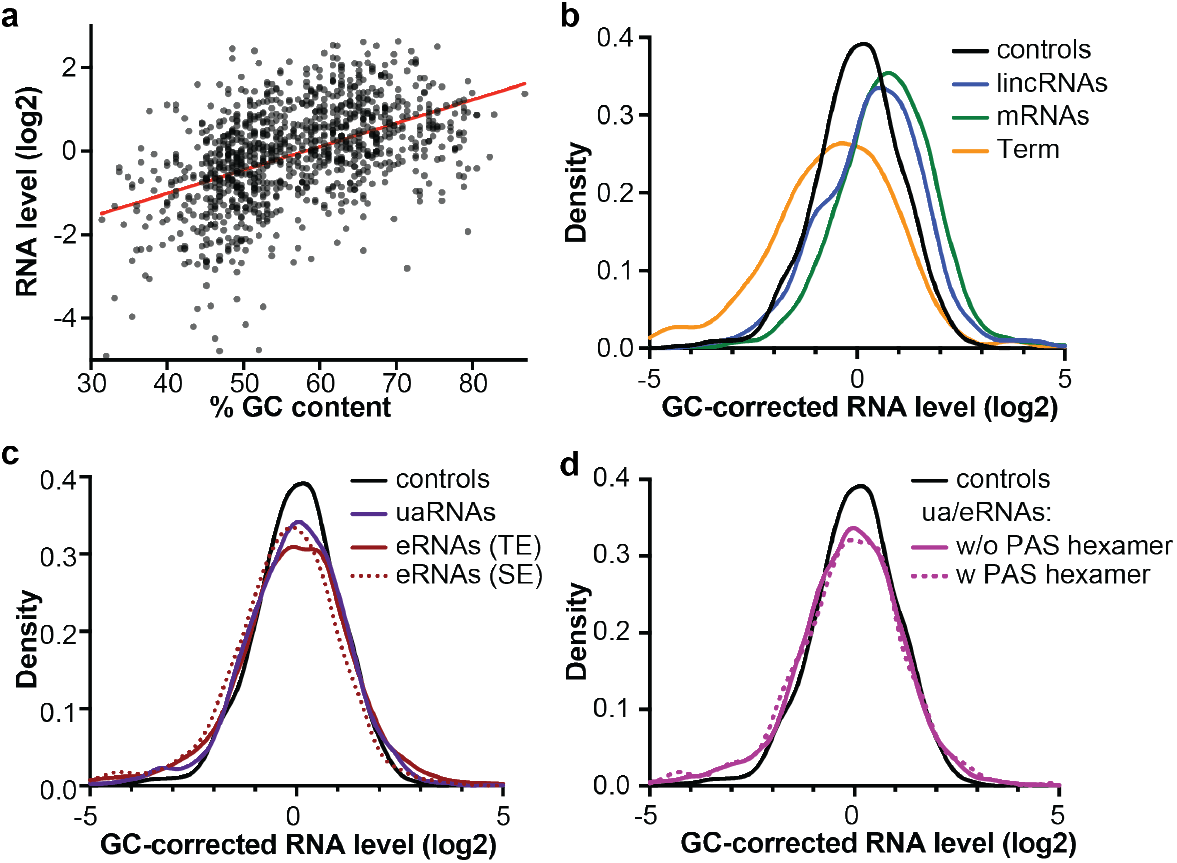
GC content inherently affects transcriptional output. **a**, Relation between GC content of synthetic control sequences and their steady-state RNA levels (*n* = 1,059). The red line is the best linear fit through the data. Pearson r = 0.44, *P* < 0.0001. **b**, Density plot of steady-state RNA abundance levels corrected for the predicted abundance based on the GC content (based on the fit line in panel a) by subtraction. Groups as in Fig. 1c. All three groups differ from the controls by Kruskal-Wallis test, *P* < 0.0001. **c**, Like b, but showing TSS-proximal regions from uaRNAs (*n* = 1,743) and eRNAs from typical enhancers (TE; *n* = 1,506) and super enhancers (SE; *n* = 600) alongside controls. Only SE eRNA regions were significantly different from the controls by Kruskal-Wallis test, *P* < 0.001. **d**, Density plot of GC-corrected steady-state RNA levels of all TSS-proximal and TSS-distal uaRNA and eRNA regions, grouped by the presence of a canonical PAS hexamer (AWTAAA, where W=A/T). Controls *n* = 1,059, without PAS *n* = 3,889, with PAS *n* = 509. The groups with and without PAS hexamer are not significantly (*P* > 0.05) different by Mann-Whitney test.

To probe to what extent GC content explained the behavior of genomic sequences in our library, a linear fit describing the relation between GC content and abundance was used to “correct” the abundance of all inserts. After correcting for GC content, mRNA and lincRNA 5’ sequences were still more abundant than predicted, and mRNA 3’ ends lower (Fig. 3b). However, the RNA levels of uaRNAs and eRNAs from typical enhancers (TEs) are indistinguishable from the controls after GC-content correction (Fig. 3c). Only eRNAs from SEs showed a small but significant shift towards lower abundance (Fig. 3c), supporting the idea that these sequences could have evolved to induce more efficient transcription termination.

Notably, our data imply that the low abundance of eRNA and uaRNA species can be explained by GC content alone, without evoking specific sequence motifs. In fact, the presence of PAS hexamers in uaRNAs and eRNAs caused no additional reduction in RNA level (Fig. 3d), demonstrating that these regions exhibit low elongation potential^5,6^ independent of PAS sequences. We conclude that most uaRNA and eRNA sequences have not evolved to contain specific motifs that minimize transcriptional output. Instead, the low GC content of the genomic regions from which these RNA species arise inherently causes RNAPII to be termination prone. Indeed, high AT content increases RNAPII pausing and backtracking^29^, and promotes termination by other RNA polymerases^30–32^, since AT-rich RNA-DNA hybrids destabilize the ternary elongation complex. In addition, AT-rich RNAs fail to form stable secondary structures that prevent RNAPII backtracking and arrest^33,34^. Thus, we propose that frequent RNAPII stalling and backtracking within AT-rich non-coding regions increases the chance of early termination, as it does downstream of a PAS at mRNA 3’ ends^35^.

### Co-transcriptionally spliced introns boost transcription

We then focused on mRNA TSS-proximal regions, which exhibited the strongest evidence for positive signals. Searches for sequence motifs that could convey these signals using MEME^36^ and Homer^37^ identified the same top hit enriched among the most abundant TSS-proximal mRNA inserts: a 5’ splice site (5’SS) motif. The 5’SS and splicing generally have been suggested to stimulate RNA production^8–17^, and the mechanisms underlying this activity are currently a subject of great interest^18^. Thus, we used INSERT-seq to systematically assess the effect of intronic sequences on RNA production. We first measured the co-transcriptional splicing efficiency of inserts in our nascent RNA screens (see methods) and found that in mRNA TSS-proximal regions, efficiently spliced inserts had the highest abundance (Fig. 4b). Further, inserts designed to contain full annotated mouse introns showed increased abundance that corresponded to splicing efficiency (Fig. 4c). This result was consistent across all screen readouts (Extended Data Fig. 4). To confirm these findings, an intron was inserted into the intron-less uaRNA reporter locus. Clonal cell lines containing at intron at this locus displayed significantly elevated RNA and protein expression as compared to intron-less controls (Fig. 4d). We note that the locus where the sequences were integrated contains no PAS hexamers, thus the stimulation caused by spliced introns does not rely on the suppression of PAS-mediated transcript cleavage or termination.

**Figure 4.**
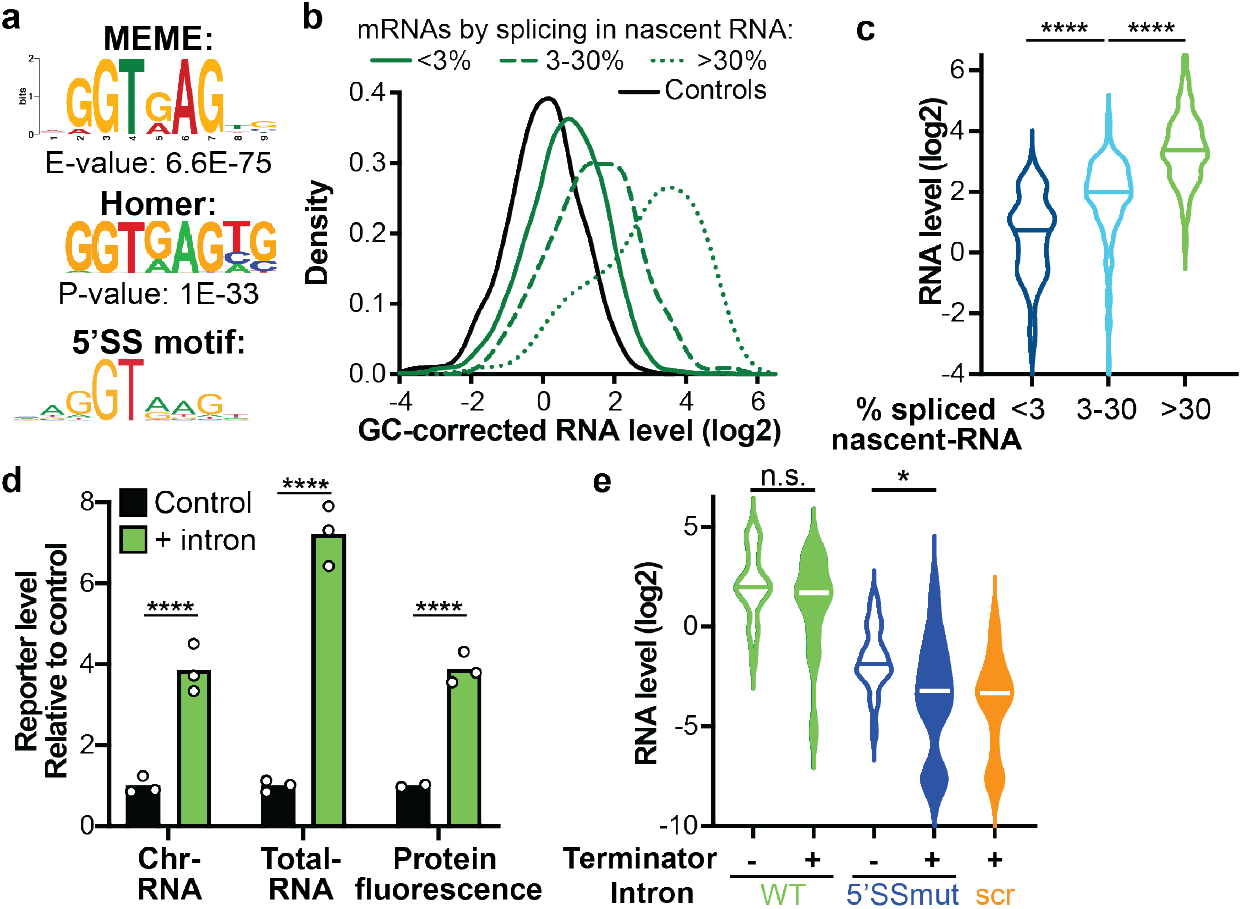
Co-transcriptionally spliced introns boost transcription. **a**, *De novo* motif most significantly enriched in the top-10% relative to the bottom-50% of TSS-proximal mRNA regions by GC-corrected steady-state RNA level. 5’ splice site (5’SS) motif from the JASPAR database (SD000.1) is shown for comparison. **b**, Density plot of GC-corrected steady-state RNA levels of TSS-proximal mRNA regions grouped by splicing efficiency (spliced / total). Controls *n* = 1059, <3% spliced *n* = 3,683, 3-30% spliced *n* = 102, >30% spliced *n* = 78. Splicing efficiency for panels b and c was calculated using nascent RNA data. All groups differ significantly from each other by Kruskal-Wallis test, *P* < 0.0001. **c**, Steady-state RNA levels of inserts containing wild-type introns (unbarcoded) grouped by splicing efficiency. <3% spliced *n* = 76, 3-30% spliced *n* = 107, >30% spliced *n* = 198, significance tested by Kruskal-Wallis test. **d**, Reporter level measured in clonal mESC lines with an intron from the mouse *Atp5a1* gene integrated upstream of the fluorescent reporter at the uaRNA locus, compared to control lines (CRISPR clones without intron integration). Effects in chromatin-associated (Chr) and steady-state (Total) RNA were determined by RT-qPCR and normalized to an internal control gene (*TBP*). Fluorescence was measured by flow cytometry. Comparisons were made using two-way ANOVA and corrected with the Šidák method. **e**, Steadystate RNA levels of variations of a subset of introns. Introns without (open) and with (filled) a strong PAS terminator^38^ replacing part of the intronic sequence were compared in otherwise wildtype (*n* = 22) and 5’SS mutant introns (*n* = 22). For reference, strong PAS terminators were inserted in a background of scrambled (scr) introns (*n* = 20). Comparisons by Mann-Whitney test. For all statistical tests in this figure: n.s. *P* > 0.05, \**P* < 0.05, \*\*\*\**P* < 0.0001.

Nonetheless, previous work indicates that the presence of a 5’SS suppresses the recognition of a downstream ‘cryptic PAS’^6–8,15–17^. To rigorously test this model, we inserted a strong 49-nt PAS-containing terminator^38^ inside a number of introns. If the terminator leads to RNA cleavage inside an insert, that will prevent amplification of the insert, leading to low abundance in the sequencing library. While addition of the terminator sequence (Fig. 4e) into wild-type, splicing-competent introns had no effect on RNA levels, mutation of the intron 5’ splice site enabled the terminator to significantly reduce transcript abundance. In fact, RNA levels dropped to the same level as that of inserts with a terminator in a scrambled sequence background (Fig. 4e). Thus, while the PAS terminator can robustly induce termination at this locus, this property is fully masked when the terminator is within a spliced intron. To our knowledge, these data are the first to demonstrate that introns can suppress the usage of not only cryptic PAS hexamers, but even strong PAS terminators.

### Distinguishing splicing-dependent and splicing-independent effects of 5’ splice site

While splicing efficiency correlated with insert abundance, we sought to establish a causal link between co-transcriptional splicing and expression using splice site (SS) mutations. For this analysis, we focused on SS mutants that abrogated splicing in a subset of efficiently spliced wild-type introns (see methods). Small (1 and 3 nt) mutations in either 5’SS or 3’SS greatly reduced RNA and protein levels (Fig. 5a, Extended Data Fig. 5a), indicating that it is the process of splicing and not the sequence composition inside introns that promotes transcription.

**Figure 5.**
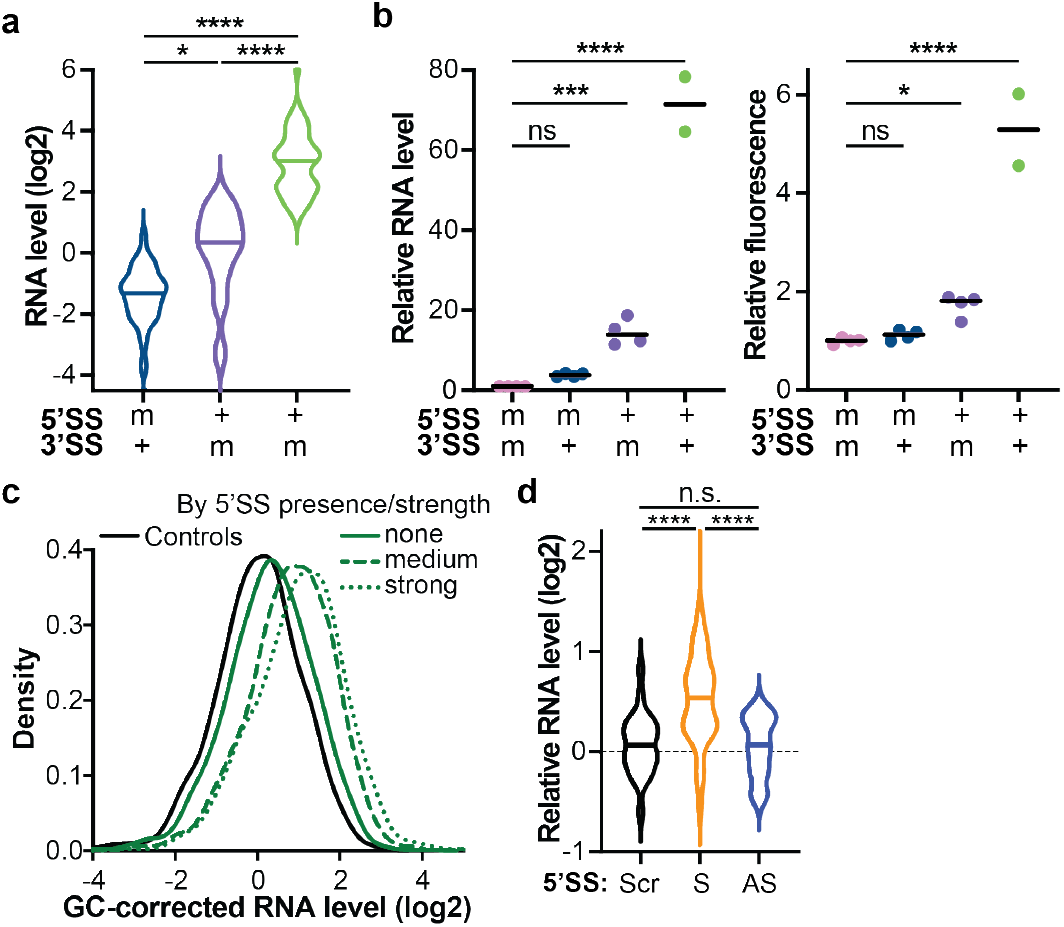
Splicing-dependent and -independent role of the 5’SS. **a**, Steady-state RNA levels of intron-containing inserts with wildtype (+) or mutant (-) splice sites. Only introns are shown of which the wild-type version was >30% spliced and mutants <3% spliced in nascent RNA. 5’SS mutants *n* = 51, 3’SS mutants *n* = 24, WT *n* = 51, comparisons by Kruskal-Wallis test. **b**, Reporter RNA level (*Tbp*-normalized RT-qPCR) and fluorescence measured by flow in different clonal cell lines containing integrated versions of an intron from the *Smc1* gene. Splice sites were either wild-type (+) or mutated (-), results were normalized to the double mutant. Comparisons made by one-way ANOVA. **c**, Density plot of GC-corrected steady-state RNA levels of unspliced TSS-proximal mRNA regions grouped by the presence and strength (MaxEnt score^39^) of a 5’SS motif (see methods). None *n* = 1,397, medium (MaxEnt 5–10) *n* = 1,783, strong (MaxEnt10+) *n* = 503. Both 5’SS-containing groups are significantly different from the inserts without a 5’SS motif (*P* < 0.0001), and from each other (*P* < 0.01) by Kruskal-Wallis test. **d**, Relative steadystate RNA levels of 10nt annotated 5’SSs with a MaxEnt score of >5, embedded into several background sequences. Only unspliced inserts (<3% spliced in nascent-RNA) were considered. Scrambled (Scr, *n* = 24) and antisense (AS, *n* = 24) versions of 5’SSs were compared to sense (S) 5’SSs (*n* = 50) by Kruskal-Wallis test. For all statistical tests in this figure: n.s. *P* > 0.05, \**P* < 0.05, \*\*\**P* < 0.001, \*\*\*\**P* < 0.0001.

Notably, 5’SS mutations had greater effects on RNA abundance than 3’SS mutations (Fig. 5a), consistent with the 5’SS having a splicing-independent role in stimulating transcription. The stronger effect of the 5’SS mutation was still observed when considering only inserts that did not contain any PAS hexamers (Extended Data Fig. 5b), demonstrating that the additional 5’SS function does not rely on suppressing PAS-mediated transcript cleavage or termination.

We confirmed the effects of both 5’SS and 3’SS mutations using clonal cell lines. In these lines, wild-type or mutant versions of a selected intron were integrated at the uaRNA locus upstream of the fluorescent reporter gene. Compared to cell lines where the insert contained mutations in both 5’SS and 3’SS, restoring just the 3’SS had no effect on RNA or protein abundance (Fig. 5b). Restoring the 5’SS alone significantly increased the reporter level, while the effect of restoring both splice sites, and thus splicing, was much larger (Fig. 5b). Importantly, restoration of either the 5’SS or 3’SS alone did not restore splicing (Extended Data Fig. 5c). This experiment supports the conclusion that the 5’SS has a splicing-independent role in stimulating transcription. Importantly, the double mutant behaves similarly to the 5’SS mutant, implying that the 3’SS only provides a positive signal in the context of a spliced intron and that the process of splicing stimulates transcription. We note that the process of splicing could strengthen the splicing-independent role of the 5’SS, or the splicing-dependent and splicing-independent stimulation could function through completely independent mechanisms.

### An autonomous role for the 5’ splice site sequence

If the 5’SS indeed promotes transcription in a splicing-independent manner, it should do so in other sequence contexts. To test this, we searched the TSS-proximal regions from mRNAs and uaRNAs for matches to the 5’SS motif consensus sequence (see methods). Considering only unspliced or very lowly spliced inserts, we find that regions containing a strong 5’SS (predicted by the MaxEntScan algorithm^39^) are more abundant than those without a 5’SS in the context of both mRNA and uaRNA/eRNA TSS-proximal regions (Fig. 5c, Extended Data Fig. 5d). This shows that a 5’SS sequence can stimulate transcription even in a context where it is not used as a splice site.

To further test whether the 5’SS sequence can function independent of sequence context, we determined if it could stimulate transcription in a random background sequence. To this end, strong 5’SS sequences (10bp) from fifty different introns were embedded in five random background sequences. The presence of a 5’SS in the sense orientation led to significant, consistent increases in expression, while the same sequences in the antisense orientation did not (Fig. 5d, Extended Data Fig. 5e). Notably, the orientation dependence points to the 5’SS sequence being recognized in the RNA, likely by the U1 snRNP. We conclude that regardless of surrounding sequence context, recognition of a 5’SS in the initially transcribed RNA by U1 snRNP promotes RNAPII elongation potential^18^.

## Discussion

In summary, using INSERT-seq we compared the effects of thousands of integrated sequences on nascent RNA, steady-state RNA and protein levels. This elucidated several novel features of transcription regulation by the initially transcribed sequence. Importantly, because all sequences were inserted at the same genomic locus, we could isolate the effects of sequence from confounding factors such as variable promoter strengths and/or chromatin contexts. Thus, the generated data sets allowed us to study causal relationships. From our results, we draw several central conclusions:

Firstly, the GC content of the initially transcribed sequence inherently affects transcriptional output. Using randomly generated sequences, we found that inserts with high GC content yield higher transcription levels and elevated RNA abundance. This likely results from high rates of RNAPII pausing and backtracking on less thermodynamically stable AT-rich sequences^33,34^. We suggest that RNAPII is particularly sensitive to this during early elongation, before all elongation factors have assembled to make RNAPII optimally processive. While it is known that GC content can influence steady state RNA accumulation^40^ and is different between coding and noncoding regions, to our knowledge GC-richness has not been previously proposed to contribute to differences in transcription levels among RNA species.

Secondly, there is no evidence that uaRNA or eRNA sequences have evolved to contain specific signals that govern RNAPII elongation. The effects of inserting uaRNA and eRNA sequences can be predicted solely based on the high AT content of intergenic regions (with some exceptions, e.g. in super enhancers). Thus, we envision RNAPII being generally termination-prone when transcribing AT-rich non-coding RNAs, rather than recognizing a specific termination signal. Importantly, this model is fully consistent with the observation that both eRNAs and uaRNAs have highly heterogeneous 3’ ends^2^.

Thirdly, we identified that mRNA 5’ ends contain positive signals, and the strongest of these are related to splicing. Our work shows that 5’SSs stimulate transcription through both splicing-dependent and -independent mechanisms. Remarkably, a 5’SS can suppress the usage of strong PAS terminators, validating and extending the telescripting model. However, we show that the effect of 5’SSs goes beyond this role, to stimulate transcription more broadly. We propose that changes in RNAPII conformation that occur upon U1 binding make it intrinsically more processive, and/or prevent association of other termination machineries, such as Integrator^2,41^ and WDR82/ZC3H4^42,43^. Providing a molecular mechanism for 5’SS-mediated transcription stimulation and identifying additional sequence elements that regulate transcription elongation will be important for future work, and we anticipate that our newly developed methodology can be leveraged towards these goals.

## Acknowledgements

We thank Kanae Sasaki for her help in optimizing the run-on protocol for screening purposes and Emily Kaye for discussions on library design. We thank Stephen Buratowski for useful discussions on the project, and Daria Shlyueva and Torben Heick Jensen for feedback on the manuscript. We are also grateful to the Flow Cytometry Facility at HMS’ Department of Immunology for their help, and to the HMS Biopolymers Facility for next-generation sequencing. This research was supported by the European Molecular Biology Organization (ALTF 531-2017 to H.V.), Human Frontier Science Program (LT000651/2018-L to H.V.), the National Institutes of Health (NIH R01 GM139960 to K.A.), startup funding from Harvard Medical School to K.A, the National Science Foundation Graduate Research Fellowship (DGE1745303 to C.A.M) and the Canadian Institutes of Health Research (Banting fellowship to B.J.E.M.).

## Author Contributions

H.V. and K.A. conceived the study and designed experiments. H.V. performed experiments and analyzed data. C.A.M. helped generate intron-containing clonal cell lines and optimize the run-on assay and knockdown conditions. B.J.E.M. and A.R.F performed ChIP-seq and TT-seq experiments. K.A. supervised the study. H.V. and K.A. wrote the manuscript with input from all co-authors.

## Competing Interests statement

K.A. is a consultant for Syros Pharmaceuticals and is on the scientific advisory board of CAMP4 Therapeutics

