## Supplementary material - figures, methods, tables for "Screening thousands of transcribed coding and non-coding regions reveals sequence determinants of RNA polymerase II elongation potential"

### Table of Contents

- Extended Data Figures 1-5
- Methods
- Supplementary Tables 1-3
- Supplementary References

### Extended Data Figures

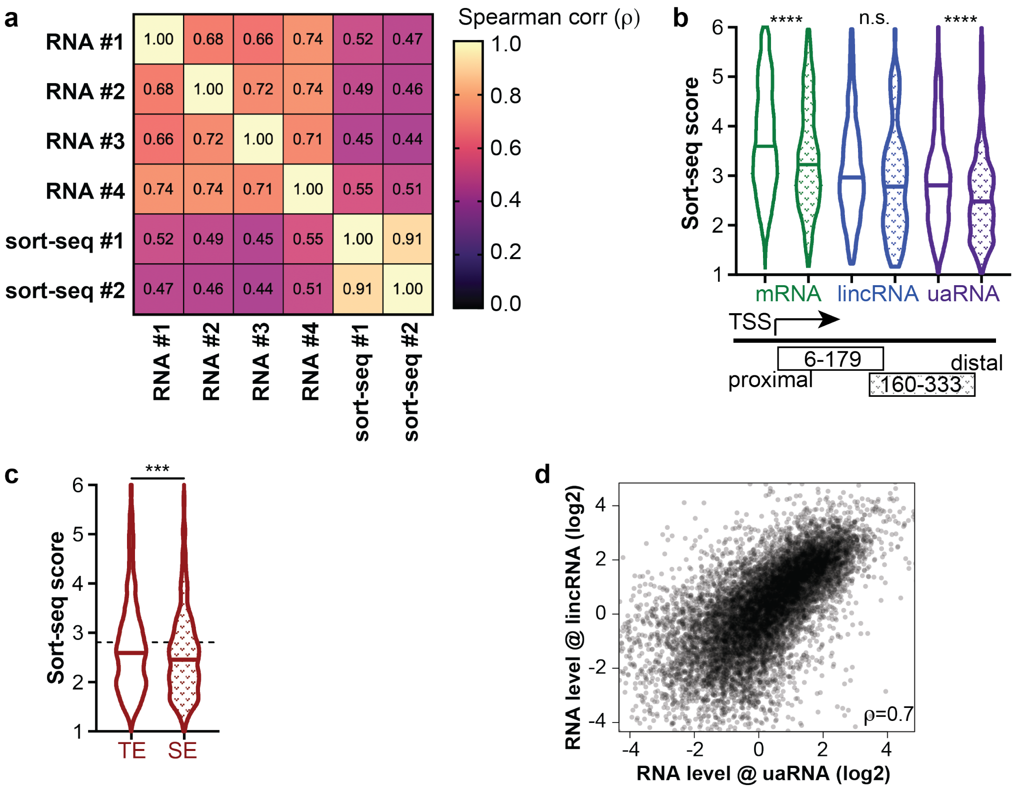


**Extended Data Figure 1. Correlations between experiments and sort-seq results.**
**a**, Spearman correlation coefficients between steady-state RNA and Sort-seq experiments, using all inserts for which data was obtained in each of the six experiments (n = 12,090). **b**, Sort-seq scores of inserts containing TSS-proximal and TSS-distal genomic regions of indicated RNA classes. Same groups as in Fig. 1e. Comparisons between proximal and distal regions by Kruskal-Wallis test (n.s. $P > 0.05$, ****$P < 0.0001$). **c**, Sort-seq scores of inserts containing TSS-proximal regions from typical enhancers (TE, $n$ = 1,506) and super enhancers (SE[1], $n$ = 600), compared by Mann-Whitney test (***$P < 0.001$). **d**, Correlation between steady-state RNA levels at the *Oct4* uaRNA locus (average of 4 replicates) and *4930461G14Rik* lincRNA locus (average of 3 replicates). Plotted are all the inserts used for Fig. 1.

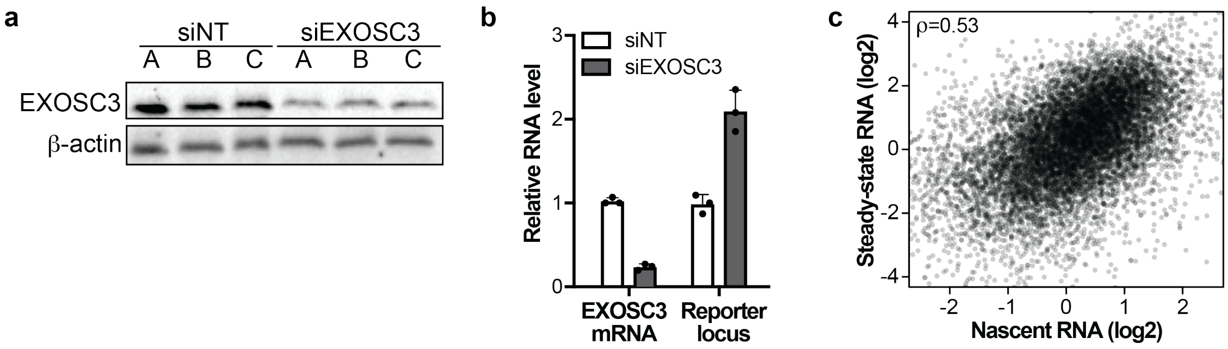


**Extended Data Figure 2. EXOSC3 knockdown validation and correlation between nascent RNA and steady-state RNA results.**

**a**, Immunoblot showing EXOSC3 protein level in control and siEXOSC3 conditions, harvested from the same experiment as the screen in Fig. 2a,b. **b**, RT-qPCR on steady-state RNA samples with which the screen was performed, showing levels of the *EXOSC3* mRNA and the reporter transcript, just downstream of the library integration site, both internally normalized to TBP. **c**, Correlation between nascent RNA (average of 2 replicates) and steady-state RNA (average of 4 replicates) levels, showing all inserts used in Fig. 1.

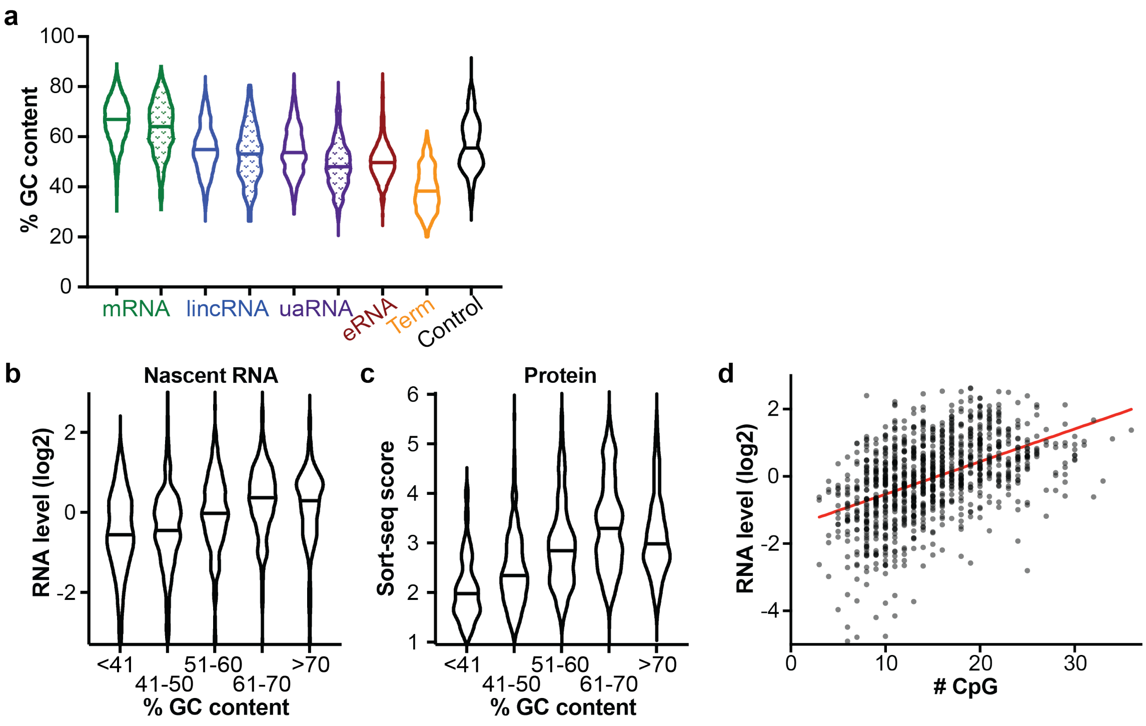


**Extended Data Figure 3. GC content in genomic region and its effect on expression.**
**a**, Distribution of GC contents in inserts of the indicated classes included in the library. Open violins show TSS-proximal regions, patterned violins show TSS-distal regions. **b**,**c**, Nascent RNA abundance (b) and sort-seq scores (c) of control sequences grouped by GC content percentage. $N = 39/281/330/292/117$ for <41/41-50/51-60/61-70/>70%, respectively. **d**, Relation between the number of CpG dinucleotides in synthetic control sequences and their steady-state RNA levels ($n$ = 1,059). The red line is the best linear fit through the data. Pearson r = 0.44, $P < 0.0001$.

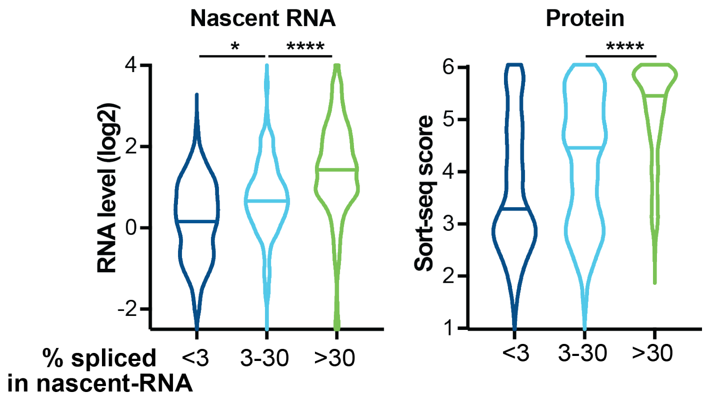


**Extended Data Figure 4. Co-transcriptionally spliced introns boost transcription and protein expression.** Nascent RNA levels (left) and Sort-seq scores (right) of inserts containing wild-type introns (unbarcoded) grouped by splicing efficiency measured using the nascent RNA screen data. <3% spliced $n$ = 76, 3-30% spliced $n$ = 107, >30% spliced $n$ = 198, significance tested by Kruskal-Wallis test. *$P < 0.05$, ****$P < 0.0001$.

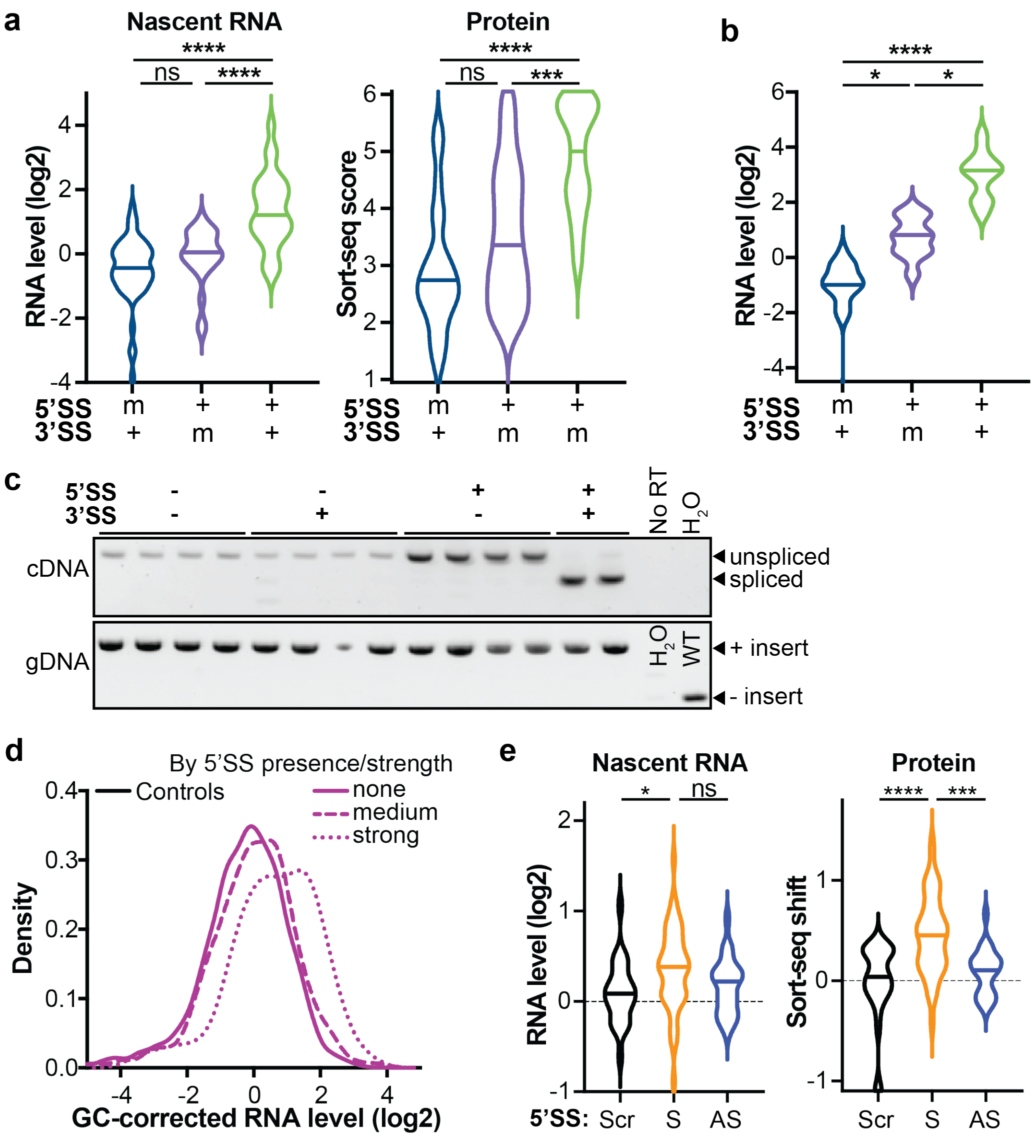


**Extended Data Figure 5. Effects of splice site mutants and 5'SS insertion in screens and clonal introns.**

**a**, Nascent RNA levels (left) and Sort-seq scores (right) of intron-containing inserts with wild-type (+) or mutant (-) splice sites. Only introns are shown of which the wild-type version was >30% spliced in nascent RNA and mutants were <3% spliced. 5'SS mutants *n* = 51, 3'SS mutants *n* = 24, WT *n* = 51, comparisons by Kruskal-Wallis test. The differences between 5'SS and 3'SS mutants was not significant in these analyses, but the pattern of the 3'SS mutants being more abundant on average was consistent with the steady-state RNA result (Fig. 5a). **b**, Steady-state RNA levels of intron-containing inserts with wild-type (+) and mutant (-) splice sites as in Fig. 5a, but showing only inserts that do not contain a PAS hexamer (any of the top-10 PASs in mouse[2]). 5'SS mutants *n* = 19, 3'SS mutants *n* = 10, WT *n* = 20, comparisons by Kruskal-Wallis test. **c**, Characterization of all clonal cell lines shown in Fig. 4c, where versions of the 14$^{th}$ intron of the Smc1 gene with wild-type (+) or mutant (-) splice sites were integrated at the *Oct4* uaRNA reporter locus. Top shows RT-PCR, bottom shows PCR on genomic DNA. All clonal lines show

**Extended Data Figure 5 (cont.)**

genomic integration of the same size in the genomic DNA, but only lines where the intron is flanked by two wild-type splice sites show evidence of splicing. Note that lanes should not be quantitatively compared to each other, as amounts of template material were not controlled. **d**, Density plot of GC-corrected steady-state RNA levels of unspliced TSS-proximal uaRNA/eRNA regions grouped by the presence and strength (MaxEnt score[3]) of a 5'SS motif (see methods). None $n = 2{,}626$, medium (MaxEnt 5–10) $n = 1{,}544$, strong (MaxEnt10+) $n = 105$. All groups are significantly different from each other ($P < 0.0001$). **e**, Relative nascent RNA levels (left) and sort-seq scores (right) of 10nt annotated 5'SSs with a MaxEnt score of >5, embedded into several background sequences. Only unspliced inserts (<3% spliced in nascent-RNA) were considered. Same groups as in Fig. 5d: scrambled (Scr, $n = 24$) and antisense (AS, $n = 24$) versions of 5'SSs were compared to sense (S) 5'SSs ($n = 50$) by Kruskal=Wallis test.
For all statistical tests in this figure: n.s. $P > 0.05$, *$P < 0.05$, ***$P < 0.001$, ****$P < 0.0001$.

### Methods

Mouse embryonic stem cell lines
All experiments were performed in the F121-9 line, a Castx129 female mouse hybrid ES cell line[4]. T2A-TagBFP2 was integrated using CRISPR-Cas9 at the location of the Oct4 stop codon by co-transfecting plasmids pHV123 and pKW09 (for all plasmids used in this study, see Supplementary Table 1, for primers see Supplementary Table 2). All transfections in this study were done using suspension transfections with Lipofectamine 2000 (Thermo Fisher). After sorting for GFP+ and TagBFP2+ cells, a clonal line was selected with integration at only the 129 allele of the Oct4 C terminus, which was validated by Sanger sequencing SNPs just outside the homology arms. In this line, mKate2 was inserted in the Oct4 uaRNA, 172bp downstream of the uaTSS, by co-transfecting pHV111 and pHV109. After sorting for mKate2+ cells, a clonal line was selected with integration at only the 129 allele. These cell lines maintained expression of the red and blue reporter proteins, as well as good mESC morphology, over many passages.
Clonal lines with an the 9th intron of the mouse *Atp5a1* gene (ENSMUST00000026495.15) inserted just upstream of the reporter, 172bp downstream of the Oct4 uaRNA TSS, were generated by co-transfecting pHV114 and a linear double-stranded repair template. The repair template was generated with two steps of PCR from a DNA fragment (ordered from Twist Bioscience) containing the intron (including 3nt each of flanking exons) and homology to the uaRNA reporter locus. The first PCR introduced more homology to the locus, the second PCR introduced phosphorothioate bonds on both ends of the amplicon (see Supplementary Table 2 for primers). After transfection, GFP+ cells were sorted, and three intron-containing clonal lines were identified by genotyping. Clones that went through the same procedure but only had a 1bp insertion at the cut site were used as control lines. Wild-type and mutant versions of the 14th intron of *Smc1a* (ENSMUST00000045312.6) with a small part of the flanking exons were integrated upstream of mKate2 by co-transfecting pHV137 with pHV163/164/165/166, sorting for GFP+ cells and genotyping individual clonal lines.
All mESC lines were cultured in KO-DMEM media (Gibco) supplemented with 15% KO Serum replacement (Gibco), GlutaMAX, penicillin-streptomycin, non-essential amino acids, beta-mercaptoethanol, 1000 U/ml LIF (Cell Guidance Systems), 1$\mu$M MEK inhibitor (PD0325901; Stemgent), and 3$\mu$M GSK3 inhibitor (CHIR99021; Stemgent).

Design of insert library
A library of 16,461 173nt sequences was designed, mostly consisting of sequences derived from specific regions in the mouse genome (mm10). The largest class (10,774 sequences) were sequences just downstream of transcription start sites as defined by start-seq data (GSE43390)[5], using GENCODE M20 (without confidence level 4/5 transcripts) to annotate TSSs while running TSScall using a count threshold of 16, search window of 250bp and joining distance of 500bp (https://github.com/lavenderca/TSScall). Using these TSSs, TSS-proximal (+6-179) and TSS-distal regions (+160-333) from different transcript classes were selected as described in Supplementary Table 3. 503 regions of annotated protein-coding transcript ends were included as described in Supplementary Table 3 as well.
High-confidence introns were defined using GENCODE M20, selected to be between 50 and 158nt in size, and filtered based on evidence of splicing in mESCs (more details in Supplementary Table 3). The 173nt sequences included in the library start 12nt upstream of the 5'SS. For a subset of introns, multiple barcoded wild-type and mutant versions were added to the

library. 1nt and 3nt mutations were made for both the 5'SS and 3'SS. A python script (https://github.com/audy/barcode-generator) was used to generate a set of 7nt barcodes with a minimum editing distance of 5 and a maximum stretch of 3. These barcodes replaced the first 7bnt of each sequence.
Shorter sequences were placed in the context of the same 5 background sequences, see Supplementary Table 3. In these backgrounds, nucleotides 10 to 19 were replaced by 10nt 5'SSs (-3 to +7). A subset of 5'SSs was also inserted in the antisense direction, or was scrambled (no GGT remained after scrambling).
As controls, a set of 1100 completely randomly generated sequences were included, which spread a range of GC contents that was comparable to the regions taken from the genome. Sequences containing AWTAAA, AGGTR or an out-of-frame ATG were excluded.

Library cloning
The 173nt sequences described above were flanked by common regions to allow for amplification of the whole library and sequencing using common Illumina sequencing primers: TTTCCCTACACGACGCTCTTCCGATCTA[$N_{173}$]TAGATCGGAAGAGCACACGTCTGAACTCC. The reverse complement of these sequences was ordered as an oligo pool from Agilent Technologies, and was amplified for 10 cycles using NEBNext Ultra II Q5 (New England Biolabs). pHV141 (Oct4 uaRNA) and pHV152 (lincRNA) were linearized using AflII and the amplified library was inserted using Gibson assembly. The resulting DNA was concentrated and cleaned up using AMPure XP beads and electroporated into E. cloni 10G ELITE electrocompetent cells (Lucigen). A small fraction of electroporated cells was plated to estimate the number of transformants (~1.5M or more), the remainder was grown up overnight in liquid LB+Amp cultures and plasmids were isolated.

Library integration
For integration at the Oct4 uaRNA allele, first a gRNA was designed that should target uaTSS+157, but overlays a SNP and thus should only target the 129 allele in the hybrid mESCs. Using TIDE[6], the efficiency of this gRNA (pHV137) was measured to be ~75% on the 129 allele and negligible on the CAST allele. The gRNA plasmid (pHV137) and library-containing plasmid pool (pHV142) were co-transfected into the clonal cell line with the integrated TagBFP2 and mKate2 reporters at a large scale. 48 hours post-transfection, 6M GFP+ cells were sorted and propagated further. Genotyping suggested that integration had occurred in ~25% of this pool of cells. During propagation, care was taken to maintain library representation by always passaging >12M cells.
For integration at the 4930461G14Rik lincRNA allele, the same approach was used, but the gRNA in pHV149 targeted TSS+102 specifically at the CAST allele. pHV149 and the library-containing plasmid pool (pHV154) were co-transfected into wild-type F121-9 cells. 48 hours post-transfection, 5.5M GFP+ cells were sorted and propagated further. Genotyping suggested that integration had occurred in ~33% of this pool of cells. During propagation, care was taken to maintain library representation by always passaging >12M cells.

RNA-based INSERT-seq screens
For steady-state RNA read-out, >5M library-containing cells was pelleted and resuspended in Trizol (Ambion). RNA was cleaned up using Direct-zol RNA miniprep columns (Zymo

Research), treated with RQ1 DNase (Promega), then cleaned up again using a total RNA purification kit (Norgen Biotek Corp).
For nascent RNA, library-containing cells were permeabilized and run on mostly as described[7], but with a few changes. For permeabilization, buffers W and P were supplemented with 1 mM EGTA, and buffer F was supplemented with 1 mM EDTA. Cells were resuspended in 500 $\mu$L Buffer W, 10 mL Buffer P was added, and this was incubated on a Nutator for 1 minutes, then 6 minutes on ice. Cells were washed once with 10mL buffer W, before being resuspended in Buffer F. Run-on was performed on 30M permeabilized cells, and in the 2X run-on reaction mix $MgCl_2$ was present at 10 mM, SUPERase-In at 0.4 U/$\mu$L and (biotin-)rNTPs are at 50 $\mu$M each. Run-on was performed at 37°C for 15 minutes, and Biotin-11-C/UTP (Perkin-Elmer) were mixed with regular rGTP and rATP (New England BioLabs). After RNA isolation using the Total RNA Purification Kit (Norgen Biotek Corp), biotinylated RNAs were immediately bound to Streptavidin M-280 magnetic beads (ThermoFisher) and binding and washing was done as described.
Purified steady-state RNA and nascent RNA still bound to the beads was reverse transcribed with Superscript IV (Thermo Fisher), using a gene-specific primer in mKate2 annealed 221nt downstream of the variable regions. cDNA from steady-state RNA was RNase-treated, cDNA from nascent RNA was removed from the beads with heat. cDNA was concentrated using 2x RNA XP beads (Beckman Coulter) to limit the number of parallel PCR reactions that had to be set up.
Sequencing libraries were generated through 17-20 PCR cycles using NEBNext Ultra II Q5 (New England Biolabs) and NEBNext Multiplex oligos, uniquely indexing each sample. Amplicons were cleaned up using 1.5x AMPure XP beads (Beckman Coulter) and concentrations were quantified using the NEBNext Library Quant Kit for Illumina, before mixing and checking the size distribution by TapeStation (Agilent). Paired-end sequencing was performed on NextSeq500, HiSeq4000 or Novaseq6000 (Illumina, 110+50 cycles or 150+150 cycles).
For normalization purposes, genomic DNA was extracted using QuickExtract (Lucigen) in parallel with RNA extraction, and sequencing libraries were generated like described for the genomic DNA made in the sort-seq protocol.
The steady-state RNA screens after EXOSC3 depletion or control condition were performed in the same way as other steady-state RNA screens. 48 hours prior to harvesting, cells were transfected with 40 nM siRNAs, using the RNAiMAX reagent (Invitrogen) in a suspension transfection. For EXOSC3 depletion, a mix of four siGENOME siRNAs was used (Catalog ID MQ-064537-01-0002, Horizon Discovery), as a control the siGENOME Non-Targeting Control siRNA #2 was used, after a comparison of multiple non-targeting siRNAs.

Sort-seq screen
Single-cell suspension of library-containing cells were made in PBS with 1% FBS and 1mM EDTA. PI (0.1$\mu$g/mL final concentration) was added an PI-negative cells were sorted into six bins based on the ratio between red (594nm, 610/20) over blue (405nm, 450/40) fluorescence using a FACSAria cell sorter (BD Biosciences). Sorting on the ratio reduces noise introduced by cell-to-cell variation in cell size. Gates were set so that the middle four bins were of equal width in the blue x red scatter plot, while the outer two bins were wider to include even cells with larger changes. Compared to control cells in which the pHV137 Cas9+gRNA plasmid had been co-transfected with a repair template that only made a point mutation to prevent re-cleavage, the

population of library-containing cells had the same median red/blue ratio, but a wider distribution.
Cells collected in each fluorescence bin, as well as unsorted cells, were pelleted and genomic DNA was extracted using QuickExtract (Lucigen). Genomic DNA was also extracted from a pellet of unsorted cells. In a first round of PCR, the Oct4 uaRNA locus was amplified from these samples with NEBNext Ultra II Q5, using primers that would not amplify remaining plasmid or aspecific integrations at other loci. Amplicons were cleaned up using 0.9X AMPure XP beads and a small fraction was used for a genotyping PCR, which showed an enrichment of insert-containing alleles in the outer bins, relative to alleles that did not integrate an insert, which were mostly found in the middle two bins.
The amplicons from the first round of PCR were used as template in a PCR with the NEBNext multiplex oligos to generate the sequencing library. This PCR and all subsequent steps were done as described for the RNA screen, with two changes: samples were amplified for ~22 cycles total between the first and second PCR step, and after pooling the sample was size-extracted from a 1% agarose gel.

Counting reads per insert

To map inserts amplified from genomic DNA, first Cutadapt v1.14[8] was used to trim off adapters and low quality ends, then bowtie2 (version 2.3.4.3)[9] was used to map against an index made of the library of 16,462 insert sequences. For bowtie2 mapping, the following parameters were set: --no-discordant --no-mixed --rdg 20,5 --rfg 20,5 --score-min L,0,-0.11. This basically disallowed gaps and allowed up to 3 mismatches on high-quality bases, with the editing distance between any two sequences in the library being >5.
For the cDNA-derived samples, first splice junctions were called with STAR (version 2.7.0)[10], using as input a concatenated fastq file with the trimmed reads of all steady-state RNA-derived libraries and a reference genome made from the full library of sequences. The following flags were used to run STAR: --outSJfilterOverhangMin 10 4 4 4 --peOverlapNbasesMin 10 --peOverlapMMp 0.08 --alignEndsType EndToEnd --outFilterMismatchNmax 5 --scoreDelOpen -10 --scoreInsOpen -10. The detected splice junctions were filtered down to only junctions on the + strand with >10 read counts and >25% of the maximum counts for other junctions detected in the same insert. Bowtie2 was ran with the same settings as above to map the cDNA libraries to a bowtie2 index that contained all original sequences, as well as the spliced versions detected by STAR.
Samtools v1.9[11] was used to output the number of reads mapped to each sequence in the 'reference genome'.

Analysis of INSERT-seq RNA data

Data of steady-state RNA, nascent RNA and corresponding genomic DNA was first normalized by setting the total of mapped reads to 1. Abundances of all unspliced and spliced versions of an insert were added up for a total abundance per insert per sample. A cutoff was applied to filter out inserts very lowly abundant in genomic DNA, for which no reliable RNA/gDNA ratio could be calculated. RNA data was first normalized to abundance in the gDNA data, and then normalized to controls, so that the median RNA/gDNA ratio of ~1100 random control sequences was 1. Per insert, a splicing efficiency was calculated by calculating the spliced/total ratio, where 'spliced' is the summed abundance of all spliced versions (0 if no spliced versions were detected). In plots, only data of inserts with complete data (i.e. nascent RNA + steady-state RNA + sort-seq, or siNT + siEXOSC3) was shown, so that the same inserts are compared throughout.

Sort-seq analysis

A cutoff was applied to filter out very lowly abundant inserts for which no reliable distribution could be calculated. To calculate a distribution over the bins for each insert, it has to be considered that the percentage of insert-containing cells can vary between bins and that sequencing yields no data on this percentage, necessitating a different method to calculate the scaling factor for each bin. After setting the total of mapped reads to 1 in each sequencing sample, a linear model was fitted to predict the composition of the unsorted sample using the composition of each of the six bins. The coefficients of this model (which fit the data well, with a Pearson correlation of >0.9 between predicted and observed) were used to scale the different bins and calculate per insert what percentage was found in each of the bins. From this the sort-seq score was calculated by assigning each bin a score of 1 through 6 and calculating a weighted average per insert.

Analysis of sequences embedded in constant background regions

Short sequences were all evaluated in the context of the same 5 background sequences. For each background, the median signal of 30 different scrambled 10-nt 5'SSs was used to normalize all sequences in that background to. For RNA assays, normalization happened by dividing by the median of scrambled 10-nt 5'SSs, for sort-seq this median was subtracted. The mean effect of an embedded sequence across all backgrounds was then calculated.

Analysis of barcoded (mutant) introns

For barcoded (versions of) introns, original introns were included in the library with three barcodes, and 1bp and 3bp mutations of the 5'SS and 3'SS with two barcodes each. Barcodes were randomly matched up with mutant versions, allowing us to determine that none of the barcodes affected the results. Since the 1bp and 3bp mutations per splice site behaved very similarly, they were considered together. As an average value for each intron version (original, 5'SS mutant, 3'SS mutant) in each assay, the median of all barcoded replicates in all experiments was used. To assess the effects of mutations, only introns of which the original version was originally spliced (>30% in nascent RNA) and mutants that reduced splicing efficiency to <3% in nascent RNA were considered.

Motif searching algorithms

To find sequence motifs that may convey positive signals, the MEME[12] and Homer[13] algorithms were used to identify motifs enriched in the top-10% versus bottom-50% of TSS-proximal mRNA regions. Searches were limited to the given strand. For MEME the ZOOPS Motif Site Distribution was expected, and motifs were 6-12 nt long. The Homer search was limited to motifs of 6, 8 or 10 nt.

Finding 5'SS motifs using MOODS and MaxEnt

MOODS v1.9.3[14] was used to find the identity and location of all sequences matching the 5'SS motif (motif SD0001.1 from the JASPAR database) using a lenient cutoff ($P < 0.05$). Only motif matches on the plus strand were considered, and the MaxEnt score[3] for each motif match was obtained. Each insert was assigned the maximum MaxEnt score of any of its 5'SS motif matches.

Since matches motif matches with a MaxEnt score of <5 did not seem to have an effect, only sequences with a MaxEnt score of >5 were considered a 5'SS sequence.

RT-qPCR on clonal cell lines

Steady-state RNA was isolated using either Trizol (Ambion) and Direct-zol columns (Zymo Research) or RNeasy columns (Qiagen). To isolate chromatin-associated RNA from intron-containing cell lines, we followed a published protocol[15], but 0.1% Triton-X-100 was added after pellets were resuspended in buffer B as well. Incubations were all performed on ice, 8 minutes in buffer A and two incubations in buffer B for 15 minutes. The remaining pellets were resuspended in Trizol, RNA was extracted using Direct-zol columns (Zymo research). All RNA was DNase treated and then purified using a total RNA purification kit (Norgen Biotek Corp). cDNA was generated using Superscript IV (Thermo Fisher) with random hexamer oligos. qPCRs were performed using a home-made SYBR mastermix on a Bio-Rad CFX384 qPCR instrument.

Plotting and statistical tests

All violin plots were made in Graphpad Prism v9.1 with smoothing set to medium. Lines of density plots were calculated in R. Unless otherwise indicated, comparisons between groups were done by Kruskal-Wallis test, and Dunn's multiple testing correction was applied to multiple comparisons within a plot.

ChIP-seq genome profile snapshots

Genomic profile snapshots were generated using with the help of the fluff software package[16]. Published H3K4me1 data[17] was downloaded and converted to FASTQ from the SRA using the SRA toolkit (accessions SRR1202461 and SRR1202462). H3K4me3 and H3K27ac ChIP-seq data were generated as follows. mESCs were cross-linked for 5 (H3K4me3) or 10 minutes (H3K27ac) with 1% formaldehyde and quenched with 0.25μM glycine. Chromatin fragmentation was performed in sonication buffer (20mM Tris pH 8.0, 2mM EDTA, 0.5mM EGTA, 0.5% SDS, 0.5mM PMSF) with a Qsonica Q800R bath sonicator at 70% amplitude pulsing with 15 second "ON" and 45 seconds "OFF" cycles for 30 (H3K4me3) or 20 (H3K27ac) minutes total sonication time. Fragment sizes were checked by agarose gel. Fragmented chromatin from 7.5M (H3K4me3) or 2.5M (H3K27ac) mESCs was used as input for immunoprecipitation. For the H3K27ac ChIP-seq, mESCs were spiked with sonicated chromatin from 0.5M S2 cells. ChIP material was diluted with 1mL of IP buffer (20mM Tris pH 8.0, 2mM EDTA, 0.5% Triton X-100, 150mM NaCl, 10% Glycerol, 5% BSA) and pre-cleared with 10μL DynaBeads Protein A (ThermoFisher) for 1 hr at 4°C. The supernatant was then further diluted with 250μL IP buffer and incubated with 16 μL H3K4me3 antibody (EpiCypher Cat#13-0028) or 10μL H3K27ac antibody (Active Motif cat#39133, lot#22618011) overnight at 4°C. Bound chromatin was isolated with 150 μL (H3K4me3) or 50μL (H3K27ac) of pre-blocked DynaBeads Protein A for 2 hrs at 4°C. Beads were washed 1x in low salt (20mM Tris pH 8.0, 2mM EDTA, 1% Triton X-100, 150mM NaCl, 0.1% SDS), 3x in high salt (20mM Tris pH 8.0, 2mM EDTA, 1% Triton X-100, 500mM NaCl, 0.1% SDS), 1x in LiCl (20mM Tris pH 8.0, 2mM EDTA, 250mM LiCl, 1% IGEPAL, 1%[w/v] sodium deoxycholate), and 2x in TE (10mM Tris pH 8.0, 0.1mM EDTA) prior to elution with 2x 250uL of elution buffer (10% SDS, 100mM sodium bicarbonate). Crosslinks were reversed by the addition of 0.2M NaCl and incubation at 65°C overnight. The IP mixture was treated with Proteinase K (Invitrogen) for 1 hr at 50°C. DNA was extracted with phenol:chloroform:isoamyl alcohol and precipitated with ethanol. Sequencing libraries were

prepared using the NEBNext Ultra II DNA Kit (New England Biolabs) as per the manufacturer's protocol. H3K4me3 and H3K27ac ChIP-seq libraries were sequenced on an Illumina NextSeq using a paired end 160 (read1:80, index:6, read2:80) and 80 cycle (read1:43, index:6, read2:42) kits respectively.
Data was mapped to mm10 (GRCm38) using Bowtie v1.2.2[18]. For H3K4me1 and H3K4me3, the following Bowtie parameters were used: -k1 -v2 -X1000 –best, and bedgraphs mapping the midpoints of deduplicated fragments in 50 bp bins were generated using custom scripts. For H3K4me1 data, fragment lengths of 150bp were assumed. For H3K27ac, the following Bowtie parameters were used: -m1 -v2 -X1000 --best. Insert sizes were filtered between 75 and 500bp using BamTools v2.4.1[19]. Duplicate reads were flagged and removed with Picard v2.8.0[20]. Bedgraphs with 50bp bins were produced using custom scripts.

TT-seq

TT-seq was performed essentially as described[21], with some minor modifications. Briefly, mESCs, treated with 0.03 % DMSO for 4 hours, were labelled with 500 μM 4-thiouridine (Sigma) for 20 minutes. Cells were counted and collected in Trizol, to which was added 5% S2 cells, labelled with 4sU for 2 hours. RNA was chloroform extracted, DNase-treated, and chloroform extracted. To 60 μg total RNA was added 2X fragmentation buffer (final concentration: 75 mM Tris Cl, pH 8.3, 112.5 mM KCl, and 4.5 mM $MgCl_2$) and heated to 95 °C for 5 minutes. Fragmentation was stopped by adding EDTA to 50 mM and placing samples on ice. RNA was ethanol precipitated and resuspended in $H_2O$. Fragment sizes were checked on the TapeStation (peak size of ~800 nt). Biotinylation reaction was performed with 0.025 mg/mL MTSEA-Biotin XX (Biotum) in reaction buffer (20% *N,N*-Dimethylformamide, 1mM EDTA, 10 mM HEPES pH 8) for 45 minutes in the dark. Labelled RNA was chloroform extracted and ethanol precipitated, and resuspended in 90 μL $H_2O$. 75 μL DynabeadsTM M-280 Streptavidin (ThermoFisher) were pre-washed with decon solution (0.1M NaOH + 50mM NaCl), 2X 100mM NaCl, 2X High Salt buffer (100mM Tris Cl pH 7.4, 10mM EDTA pH 8, 1M NaCl, 0.05% (v/v) Tween 20), and resuspended in High salt buffer. Labelled RNA was denatured by heating to 65 °C for 5 minutes and placing on ice for 2 minutes. 10 μL High salt buffer was added to labelled RNA. Prewashed beads were placed on magnet, supernatant discarded, and then beads were resuspended in the labelled RNA. Beads and RNA were rotated for 30 minutes in the dark. Beads were then washed 4X for 1 minute with high salt buffer. Labelled RNA was eluted 2X with 100 mM DTT (1,4-Dithiothreitol). RNA was cleaned up with microelute columns (Norgen). Sequencing libraries were prepared from 250 ng of labelled RNA using the TruSeq Stranded Total RNA sequencing kit with RiboZero rRNA depletion, shortening the fragmentation step to 3 minutes and using 8 cycles of PCR amplification.
Reads were trimmed for a minimum quality score of 20 using a custom script (https://github.com/AdelmanLab/NIH_scripts/tree/main/trim_and_filter_PE), adapter sequences were removed using cutadapt version 1.14[8]. Reads were first mapped to dm6 using bowtie[18], before aligning to mm10 using STAR (version 2.7.3a)[10]. Strand-specific bedgraphs were generated from deduplicated BAM files using STAR.

### Supplementary Tables

#### Supplementary Table 1: plasmids used in this study

| Name | Description | Generation |
|---|---|---|
| pHV118 | T2A-mTurqoise2 flanked by homology arms for Oct4 C-terminal fusion. Only used for cloning. | Assembled by Gibson, using pUC19 digested with EcoRI and HindIII, and a gBlock containing GSG-T2A-VAT-mTurqoise2 flanked by homology arms for integration at the Oct4 C terminus. |
| pHV123 | T2A-TagBFP2 flanked by homology arms for Oct4 C-terminal fusion. | Assembled by Gibson, using pHV118 digested with NcoI and AccI, and TagBFP2 amplified from Addgene75178 with an overhang introduced in the primers. |
| pHV111 | mKate2 flanked by homology arms for cut at Oct4 uaTSS+172. | gBlock containing mKate2 and SV40 terminator with 500bp homology arms for cut at Oct4 uaTSS+172 digested with EcoRI and PstI was ligated into pUC19 backbone, also digested with EcoRI and PstI. |
| pKW09 | SpCas9(BB)-2A-eGFP and gRNA targeting just downstream of Oct4 stop codon | Oligos were annealed and phosphorylated, then ligated into BbsI-digested pX458 (Addgene 48138). |
| pHV109 | SpCas9(BB)-2A-eGFP and gRNA targeting Oct4 uaTSS + 172bp. | Oligos were annealed and phosphorylated, then ligated into BbsI-digested pX458 (Addgene 48138). |
| pHV141 | Recipient plasmid for library, has homology arms flanking Oct4 uaTSS+157 cut site on 129 allele with mKate2 reporter inserted at uaTSS+172, introduced AflII sites and sequencing adapters. | Assembled in a 3-piece Gibson with a backbone from pUC19 digested with EcoRI and HindIII, and two amplicons introducing 500bp homology arms and AflII restriction sites. Left and right homology arms were amplified from hybrid genome and plasmid pHV111, respectively, for primers see Supplementary Table 2. |
| pHV114 | SpCas9(BB)-2A-eGFP and gRNA targeting just upstream of mKate2 in reporter line. | Oligos were annealed and phosphorylated, then ligated into BbsI-digested pX458 (Addgene 48138). |
| pHV137 | eSpCas9(1.1)-T2A-eGFP and gRNA targeting Oct4 uaRNA +157 (B6/129 allele, not CAST) | Oligos were annealed and phosphorylated, then ligated into BbsI-digested Addgene79145. |
| pHV163 | pHV141 + WT SMC1a 14th intron | Assembled by Gibson, using pHV141 digested with AflII and an amplicon generated by amplifying the Smc1a 14th intron (see primer list). |
| pHV164 | pHV141 + 5'SSmut SMC1a 14th intron | Assembled by Gibson, using pHV141 digested with AflII and an amplicon generated by amplifying the Smc1a 14th intron (see primer list). |
| pHV165 | pHV141 + 3'SSmut SMC1a 14th intron | Assembled by Gibson, using pHV141 digested with AflII and an amplicon generated by amplifying the Smc1a 14th intron (see primer list). |
| pHV166 | pHV141 + 5/3'SS double mut SMC1a 14th intron | Assembled by Gibson, using pHV141 digested with AflII and an amplicon generated by amplifying the Smc1a 14th intron (see primer list). |
| pHV142 | Library inserted into pHV141 | Assembled by Gibson, using pHV141 digested with AflII and the library amplified using Inserts_fw/rv for 10 cycles. |
| pHV149 | eSpCas9(1.1)-T2A-eGFP and gRNA targeting 4930461G14Rik TSS+101 (CAST allele) | Oligos were annealed and phosphorylated, then ligated into BbsI-digested Addgene79145. |
| pHV152 | Recipient plasmid for library, has homology arms flanking 14Rik TSS+101 cut site on CAST allele, introduced AflII sites and sequencing adapters. | Assembled in a 3-piece Gibson with a backbone from pUC19 digested with EcoRI and HindIII, and two amplicons introducing 500bp homology arms and AflII restriction sites. Left and right homology arms were amplified from hybrid genome, for primers see Supplementary Table 2. |
| pHV154 | Library inserted into pHV152 | Assembled by Gibson, using pHV152 digested with AflII and the library amplified using Inserts_fw/rv for 10 cycles. |

### Supplementary Table 2: Primers used in this study

| Name | Use | Sequence |
|---|---|---|
| TagBFP2_Gibson_Fw | To amplify mTagBFP2 for Gibson | CATGCGGTGACGTGGAGGAGAATCCCGGCCCTGTCGCCACCATGAGCGAGCTGA |
| TagBFP2_Gibson_Rv | | GTAAAAGAATTTAACCCCAAAGCTCCAGGTTCTCTTGTTACCTCCCTTGCCTTGGCTCACAGCATCCCCTTAATTAAGCTTGTGCCCCAG |
| sg_Oct4_Cterm_fw | Cloning gRNA against Oct4 C terminus into plasmid (pHV123) | CACCGACTGAGGCACCAGCCCTCCC |
| sg_Oct4_Cterm_rv | | AAACGGGAGGGCTGGTGCCTCAGT |
| sg_Oct4_uaTSS+172bp_Fw | Cloning gRNA against Oct4 uaRNA region into plasmid (pKW09) | CACCGGCTGTAAGGACAGGCCGAGA |
| sg_Oct4_uaTSS+172bp_Rv | | AAACTCTCGGCCTGTCCTTACAGCC |
| sg_mKate2_2_Fw | Cloning gRNA against region upstream of mKate2 into plasmid (pHV114) | CACCGTGGTGGCGACCGGTGGATCC |
| sg_mKate2_2_Rv | | AAACGGATCCACCGGTCGCCACCAC |
| *not a primer* but dsDNA fragment | Starting point of the Atp5a1 intron repair template | TGGGTCACCGGACACCTCACAAACCAGTTGCTCGGATGCCCCATCGCACCACAAAGCCTGTTGGCACTGCACCCTCTCTAGACCGGGAcaCAGgtaagttttcctctctttgacacaggagtttgttactagcttaaaatcttaaatgtaggacctttgtcttatactgttgtgtggcattaagtcatggtgacactaacttgtgtgctcagGTGgtACCGGTCGCCACCATGGTGAGCGAGCTGATTAAGGAGAACATGCACATGAAGCTGTACATGGAGGGCACCGTGAACAACCACCACTTC |
| uaRNA_mKate2_morehomology_Fw | Amplifying Atp5a1 intron repair template while introducing more homology to the locus | gtgggtggaggagcagagctgtgggggtgggagaaactgaggcgagcgctatctgcctgtgtcttccagacggaggttggggacgtctggacaggacaacccttaggacgggaccccaggaggccttcattttcaaccttcaaggtcctctcacccctgcctTGGGTCACCGGACACCTCAC |
| uaRNA_mKate2_morehomology_Rv | | agctggtagccaggatgtcgaaggcgaaggggagagggccgccctcgaccgccttgattctcatggtctgggtgccctcgtagggcttgccttcgccctcggatgtgcacttGAAGTGGTGGTTGTTCACGGTGC |
| uaRNA_mKate2_amplify_Fw | Amplifying Atp5a1 intron repair template while introducing phosphorothioate bonds | G*T*GGGTGGAGGAGCAGAGCT |
| uaRNA_mKate2_amplify_Rv | | A*G*CTGGTAGCCAGGATGTCGA |
| sg_Oct4_uaTSS+157-mK_Fw | Cloning gRNA against Oct4 uaTSS+157 (129 allele) into plasmid (pHV137) | CACCAGAGAGGGTGCAGTGCCAAC |
| sg_Oct4_uaTSS+157-mK_Rv | | AAACGTTGGCACTGCACCCTCTCT |
| pHV141_left_fw | Amplify Oct4 homology arm to use in Gibson (for pHV141) | cacaggaaacagctatgaccatgattacgccaCCTCTGAGCCTGGTCCGATTCC |
| pHV141_left_rv | | GCTCTTCCTTAAGTGTCTTAAGGAAGAGCGTCGTGTAGGGAAAGAGTGTAACAGGCTTTGTGGTGCGATGG |
| pHV141_right_fw | Amplify Oct4 uaRNA + mKate2 homology arm to use in Gibson (for pHV141) | TCCTTAAGACACTTAAGGAAGAGCACACGTCTGAACTCCAGTCACGCACTGCACCCTCTCTAGACC |
| pHV141_right_rv | | CAGTCACGACGTTGTAAAACGACGGCCAGTGAATTGGTACAGGGTCTCGGTGGAGG |
| SMC1_intron14_WT_Fw_wGibsonoverhang | Cloning pHV163/pHV165 | ACACTCTTTCCCTACACGACGCTCTTCCGATCTACCAGgttagtaggggggtgggcatg |
| SMC1_intron14_WT_Rv_wGibsonoverhang | Cloning pHV163/pHV164 | GACTGGAGTTCAGACGTGTGCTCTTCCGATCTCCACctgagggaacagtcagggaagg |

| | | |
|---|---|---|
| SMC1_intron14_5ssmut_Fw_wGibsonoverhang | Cloning pHV164/pHV166 | ACACTCTTTCCCTACACGACGCTCTTCCGATCTACCACtgtagtaggggg tgggcatg |
| SMC1_intron14_3ssmut_Rv_wGibsonoverhang | Cloning pHV165/pHV166 | GACTGGAGTTCAGACGTGTGCTCTTCCGATCTCCACagcagggaaca gtcagggaagg |
| Inserts_fw | To amplify oligo pool and introduce more homology to recipient plasmid (pHV142/pHV154) | GTGACTGGAGTTCAGACGTGTGCTCTTCC |
| Inserts_rv | | ACACTCTTTCCCTACACGACGCTCTTCC |
| sg_4930461G14Rik_TSS+101_Fw CAST | Cloning gRNA against 4930461G14Rik TSS+101 (CAST allele) into plasmid (pHV149) | CACCATCCTAAAGATCTGAACAGT |
| sg_4930461G14Rik_TSS+101_Rv CAST | | AAACACTGTTCAGATCTTTAGGAT |
| pHV152_left_fw | Amplify Oct4 homology arm to use in Gibson (for pHV141) | CACAGGAAACAGCTATGACCATGATTACGCCATTCCAGCATGCTG GGGTCG |
| pHV152_left_rv | | GCTCTTCCTTAAGTGTCTTAAGGAAGAGCGTCGTGTAGGGAAAGA GTGTTGTTCAGATCTTTAGGATAGCCCT |
| pHV152_right_fw | Amplify Oct4 uaRNA + mKate2 homology arm to use in Gibson (for pHV141) | TCCTTAAGACACTTAAGGAAGAGCACACGTCTGAACTCCAGTCAC AGTGGGGACAGAATGAAGAGAG |
| pHV152_right_rv | | CAGTCACGACGTTGTAAAACGACGGCCAGTGAATTTCTCATGACT GGCTGGGCTC |
| mKate2_RT | Gene-specific RT primer within mKate2 | CCCTCGACCGCCTTGATTCTC |
| Oct4_uaTSS+113_Fw | First PCR round: amplify Oct4 uaRNA + mKate2 genomic locus | CACAAACCAGTTGCTCGGATGC |
| mKate2_Rv | | CACGAGCTTCAGGGCCATGTC |
| mKate2_genotyping_Rv | Genotyping to determine the % +/- insert alleles in each bin, together with Oct4_uaTSS+113_Fw | GTCTTCGTATGTGGTGACTCTCTCC |

### Supplementary Table 3: description of insert sequences in the library used for this study

| **Regions close to TSSs** | Criteria for TSS selection, of which downstream sequences were used | Nr in library | Region (distance to TSS) |
|---|---|---|---|
| mRNA TSS-proximal regions | TSSs that were associated with a protein-coding transcript, were the strongest TSS within their cluster, were focused, and PRO-seq reads could be detected downstream. | 4078 | 6-179, or shifted up/down by 1bp to shift annotated start codon, or first ATG if there is no annotated start in the region, in frame with mKate2. |
| mRNA TSS-distal regions | Random subset of TSSs selected for mRNA TSS-proximal regions | 1020 | 160-333, or shifted up/down by 1bp to shift annotated start codon, or first ATG if there is no annotated start in the region, in frame with mKate2. |
| lincRNA TSS-proximal regions | TSSs that were associated with a lincRNA, but not any protein-coding transcripts. 2/3rds met criteria on TSS focus and PRO-seq signal were included, for the other 1/3rd those criteria were not used. | 388 | 6-179, or shifted up/down by 1bp to shift the first ATG in frame with mKate2. |
| lincRNA TSS-distal regions | Random subset of TSSs selected for lincRNA TSS-proximal regions | 105 | 160-333, or shifted up/down by 1bp to shift the first ATG in frame with mKate2. |
| uaRNA TSS-proximal regions | TSSs located upstream and antisense to the 4200 selected mRNA TSSs, not associated with protein-coding transcripts. | 1914 | 6-179, or shifted up/down by 1bp to shift the first ATG in frame with mKate2. |
| uaRNA TSS-distal regions | Random subset of TSSs selected for uaRNA TSS-proximal regions | 638 | 160-333, or shifted up/down by 1bp to shift the first ATG in frame with mKate2. |
| Traditional enhancer TSS-proximal regions | Unannotated TSSs that were the highest in a cluster of multiple TSSs, marked with H3K7ac, and more than 1kb away from a gene model. Not overlapping SuperEnhancer regions[1]. | 1700 | 6-179, or shifted up/down by 1bp to shift the first ATG in frame with mKate2. |
| Super Enhancer TSS-proximal regions | 134 unannotated TSSs that were the highest in a cluster of multiple TSSs, marked with H3K7ac, and more than 1kb away from a gene model, which do overlap SuperEnhancer regions[1]. In addition, 549 more TSSs that were found within SuperEnhancers and that did not have to be the highest in their cluster. | 683 | 6-179, or shifted up/down by 1bp to shift the first ATG in frame with mKate2. |

| **Regions around TESs** | Criteria for selection | Nr in library | Region (relative to PAS at transcript end) |
|---|---|---|---|
| mRNA terminators | mRNA transcription ends from annotation, which showed a typical PRO-seq pattern where signal dips at a cleavage site and increases downstream, with a canonical PAS hexamer (AAUAAA) within 87bp. | 503 | -86 to +87 around PAS (or center of PASs) |

| **Introns** | Criteria for selection, and how seqs were modified | Nr in library (bc * seqs) | Region (relative to 5'SS) |
|---|---|---|---|
| General criteria for all: | Introns between 50 and 158bp, from transcripts that have the highest confidence in their gene in GENCODE M20, downstream of TSSs observed using start-seq and spliced out in >2/3 of annotated transcripts that contain the region. RPKM of <0.33 and <0.1 compared to the median | | |

| | | | |
|---|---|---|---|
| | of exons in the same transcript, for 4sU-seq and RNA-seq in mESCs, respectively. | | |
| First introns | Introns matching general criteria, that are the first intron in a transcript, and where the 3'SS is <750nt from the TSS. | 298 | 12bp upstream exon to 5'SS + 161bp (introns are 50-158bp, so minimally 3bp of downstream exon). |
| Non-first introns | Introns matching general criteria, that are not the first intron in a transcript (64 w/o any top-10 PAS hexamer, 60 with AAUAAA hexamer inside intron) | 124 | 12bp upstream exon to 5'SS + 161bp (introns are 50-158bp, so minimally 3bp of downstream exon). |
| Barcoded original introns | Same set as non-first introns. First 7bp (all part of the upstream exon) replaced by barcode. | 3*120 = 360 | 7bp barcode, then 5bp of upstream exon to 5'SS + 161bp (introns are 50-158bp, so minimally 3bp of downstream exon). |
| Barcoded 5'SS mut 1bp | Same set as non-first introns, first nucleotide of intron (usually G) mutated to C. First 7bp (all part of the upstream exon) replaced by barcode. | 2*120 = 240 | 7bp barcode, then 5bp of upstream exon to 5'SS + 161bp (introns are 50-158bp, so minimally 3bp of downstream exon). |
| Barcoded 5'SS mut 3bp | Same set as non-first introns, nucleotides -1 to +2 (usually GGT) mutated to CTG. First 7bp (all part of the upstream exon) replaced by barcode. | 2*120 = 240 | 7bp barcode, then 5bp of upstream exon to 5'SS + 161bp (introns are 50-158bp, so minimally 3bp of downstream exon). |
| Barcoded 3'SS mut 1bp | Same set as non-first introns, last nucleotide of intron (usually G) mutated to C. First 7bp (all part of the upstream exon) replaced by barcode. | 2*120 = 240 | 7bp barcode, then 5bp of upstream exon to 5'SS + 161bp (introns are 50-158bp, so minimally 3bp of downstream exon). |
| Barcoded 3'SS mut 3bp | Same set as non-first introns, last three nucleotides (usually CAG) mutated to GCT. First 7bp (all part of the upstream exon) replaced by barcode. | 2*120 = 240 | 7bp barcode, then 5bp of upstream exon to 5'SS + 161bp (introns are 50-158bp, so minimally 3bp of downstream exon). |
| Barcoded introns with PAS terminator | Same as PAS-hexamer-containing non-first intron with hexamer >58nt of the 3' splice site. PAS hexamer + downstream region (44-48nt total) were replaced by 49nt PAS terminator[22] and 1-5nt of the downstream exon were trimmed off. | 3*24 = 72 | 7bp barcode, then 5bp of upstream exon to 5'SS + 160/164bp, with 49nt PAS terminator inside |
| Barcoded mutant introns with PAS terminator | Same as Barcoded introns with PAS terminator, but 3bp mutation was made at 5'SS (>CTG) | 3*24 = 72 | 7bp barcode, then 5bp of upstream exon to 5'SS + 160/164bp, with 49nt PAS terminator inside |
| Scrambled introns with PAS terminator | Same as Barcoded introns with PAS terminator, but region around 49nt PAS terminator was scrambled. | 24 | Scrambled versions of intron/exon regions flanking 49nt PAS terminators |

| **Short sequences in backgrounds** | Criteria for selection | Nr in library (bc * seqs) | Region (relative to 5'SS) |
|---|---|---|---|
| Criteria for background sequences: | Randomly generated, but not containing a PAS hexamer, 5'SSs, ATGs, in-frame stop codons, or mononucleotide repeats of 5nt or longer. | | |
| 10nt 5'SS in sense orientation | 5'SSs of subset of introns that were barcoded/mutated, each in 5 different backgrounds. | 5*60 = 300 | 10bp 5'SS (-3 to +7), replacing bp 10-19 in background sequences |

| | | | |
|---|---|---|---|
| 10nt 5'SS in antisense orientation | Reverse complement of a subset of the sense 5'SSs, inserted in same location in same backgrounds. | 5*30 = 150 | 10bp 5'SS (-3 to +7), rev compl, replacing bp 10-19 in background sequences |
| 10nt 5'SS scrambled | Same subset of 5'SSs as for antisense orientation, but for each slice site the nucleotide order was randomized until no GGT remained. | 5*30 = 150 | 10bp 5'SS (-3 to +7), scrambled, replacing bp 10-19 in background sequences |

| **Control sequences** | Description | Nr in library |
|---|---|---|
| Randomized controls | Randomly generated using distribution of GC content comparable to different genomic regions, sequences containting PAS, 5'SS or out-of-frame ATG were excluded. | 1100 |
